# Synonymous coding significantly affects the domain swapping propensity of myoglobin

**DOI:** 10.64898/2026.04.02.716112

**Authors:** Shlomit Dor, Ailie Marx

## Abstract

Co-translational folding is a critical, yet poorly understood, aspect of protein biogenesis due to its transient, heterogeneous, and experimentally inaccessible nature. Using a myoglobin variant engineered towards increased domain swapping, we show that stable dimers formed during heterologous E. Coli expression revert to the monomeric state following denaturation - renaturation and that domain swapping propensity is significantly affected by synonymous coding. Wider implications for the role of synonymous coding in aggregation and disease are discussed.

## Main Text

Although some proteins can fold outside the cell, as famously demonstrated by Anfinsen’s Nobel Prize winning experiments (Anfisen et al 1961 and Anfisen 1973), this is not how the process occurs naturally. Within the cell, proteins are produced by ribosomal translation of the mRNA genetic instructions into a nascent protein chain. As amino acids are sequentially added to the growing nascent chain, it transverses the ribosome exit tunnel before entering the cytosol where it folds into a three-dimensional protein structure. The vectorial emergence of the nascent chain from the ribosome enables co-translational folding, the progressive acquisition of structure by the region of the nascent chain emerging from the ribosomal exit tunnel whilst downstream regions are still being translated (Holtkamp et al 2015). Ribosome elongation rates, influenced by synonymous codon choice, can modulate the timing of secondary- and tertiary-structure formation, alter the exposure of folding intermediates, and affect interactions with chaperones (Walsh et al., 2020; Moss et al., 2024). Studies have also shown that synonymous variation can alter protein activity and substrate specificity without changing amino-acid sequence (Kimchi-Sarfaty et al., 2007), as well as cellular fitness and proteostasis, largely through changes in translation kinetics rather than protein abundance (Walsh et al., 2020; Shen et al., 2022). Despite its importance, co-translational folding remains challenging to study experimentally, as many of its key intermediates are transient, heterogeneous, and difficult to capture using traditional structural or biochemical approaches.

Identifying robust, experimentally tractable readouts of co-translational folding therefore remains a major challenge. In this context, proteins capable of domain swapping provide a potentially sensitive system in which subtle perturbations to folding kinetics can be amplified into discrete, stable and measurable structural outcomes. The domain swapping mechanism sees two or more protein molecules form a thermodynamically stable dimer or higher oligomer by exchanging identical structural elements, around a hinge region, (Mascarenhas et al 2017, Rousseau et al., 2011; Bennett et al., 2006; Liu et al., 2001). Previous studies have shown that refolding stable domain swapped species from the unfolded states can preferentially yield monomers (Rousseau et al. 2012), an observation suggesting that domain swapping is a kinetically accessible folding product, and as such, potentially sensitive to translation rhythm and synonymous coding.

Horse heart myoglobin has been known for decades to form a small fraction of domain swapped species *in vivo* (Van den Oord et al 1969, Figure S1.a). X-ray crystal structures of the domain swapped form (Nagao et al 2012) show that domain swapping results from an extension of the loop region between helix E and helix F into a continuous alpha helical structure (Figure 1a). It has been shown that the portion of the domain swapped protein can be greatly increased by mutation of the hinge loop residues to amino acids with higher helical propensity (Nagao et al 2020, Xie et al 2021), including the triple alanine mutant that we henceforth refer to as the AAA variant. To assess the effect of synonymous coding on domain swapping propensity, we first characterized the suitability of AAA myoglobin as a partially domain swapping system where measurable amounts of both monomer and dimer are formed during cellular translation. The oligomeric populations of AAA myoglobin, expressed in *E. coli*, were observed by SDS PAGE (Figure 1c) and assessed by size exclusion chromatography (Figure 1b) demonstrating that the dimer: monomer ratio for the AAA mutant is quantifiable and reproducible. Further analysis demonstrated that both monomeric and dimeric protein products remain stable for at least 7 days at 4 degrees Celsius (Figure 1b). That the dimeric species is not even under denaturing SDS PAGE conditions further highlights the stability of the dimeric species (Figure 1c). We further found that unfolding – refolding each of the individual oligomeric populations leads to a predominantly monomeric state (Figure 1d) suggesting that the domain swapped state of AAA myoglobin is kinetically trapped.

**Figure 1.**
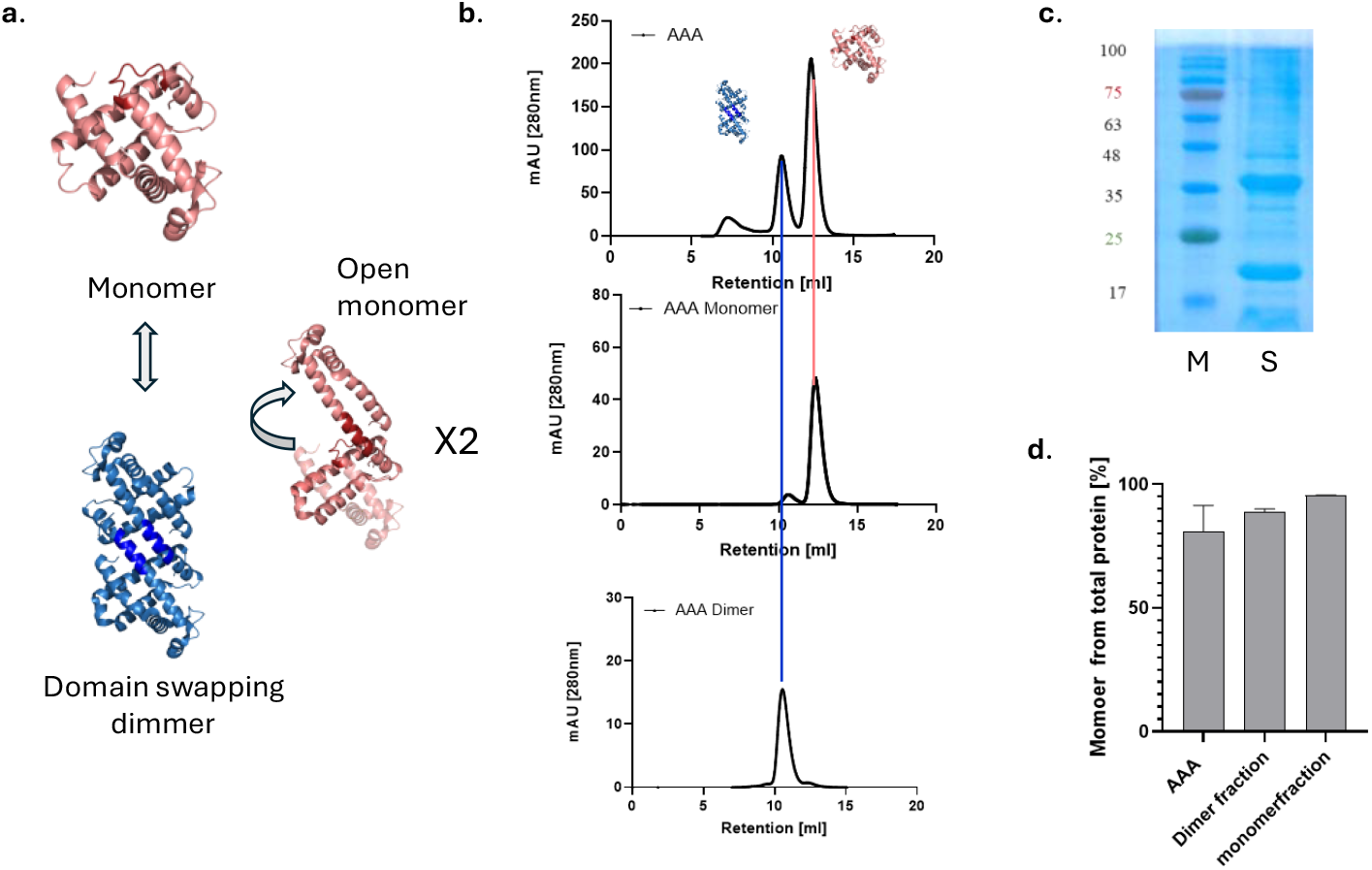
AAA myoglobin forms stable, kinetically trapped, domain swapped dimers. (a) The AAA variant of myoglobin can form either a monomer (pink) or a domain swapped (blue) conformation. (b) These populations are fully separable by size exclusion chromatography; both the monomeric and dimeric peaks remain stable after storage at 4 °C for one week. (c) 15% SDS PAGE showing the monomeric (∼19KD) and dimeric (∼40KD) species of AAA myoglobin (lane 1 (M); marker and lane 2 (S); sample) (d) Refolding profile of the AAA fractions following denaturation in 8 M urea.

To explore the hypothesis that synonymous coding can alter the domain swapping propensity of AAA myoglobin, the sequence before and after the hinge, individually, was mutated to common (NTD & CTD Common) or rare codons (NTD & CTD Rare). The exact nature of the synonymous mutations, consisting of between seventeen and twenty four synonymous alternations per construct, is shown in Figure S2 and S3. Analysis of the domain swapping propensity of the synonymous mutants showed that rare coding, both before and after the hinge region, greatly increased the dimer: monomer ratio in AAA myoglobin. A significant increase of 20% (**p<0.0049) was observed in the case of rare recoding after the hinge region (CTD Rare; Figure 2a, 2b). On the contrary, mutation to common codons, either before or after the hinge, had no effect on the dimer: monomer ratio. Although the increased domain swapping propensity was associated with increased (approximately doubled) soluble protein yield of the CTD Rare variant (*p< 0.0288 Figure 2c), there was no detectable difference in gene expression patterns between the synonymously coded variants (Figure 2d). This might suggest that increased domain swapping in myoglobin AAA prevents loss of soluble protein either by aggregation or degradation. Given that Figure S4 shows that there is no appreciable accumulation of insoluble protein in any construct as observed on SDS-PAGE, the apparently increased yield in soluble protein when rare synonymous codon mutations are introduced may be an artifact of higher order oligomerization. Notably, a higher-order oligomeric state was present in all chromatograms, though it is reduced in the rare-coded constructs. Future studies should characterize this oligomeric state and employ alternate methods for calibrating concentration.

**Figure 2.**
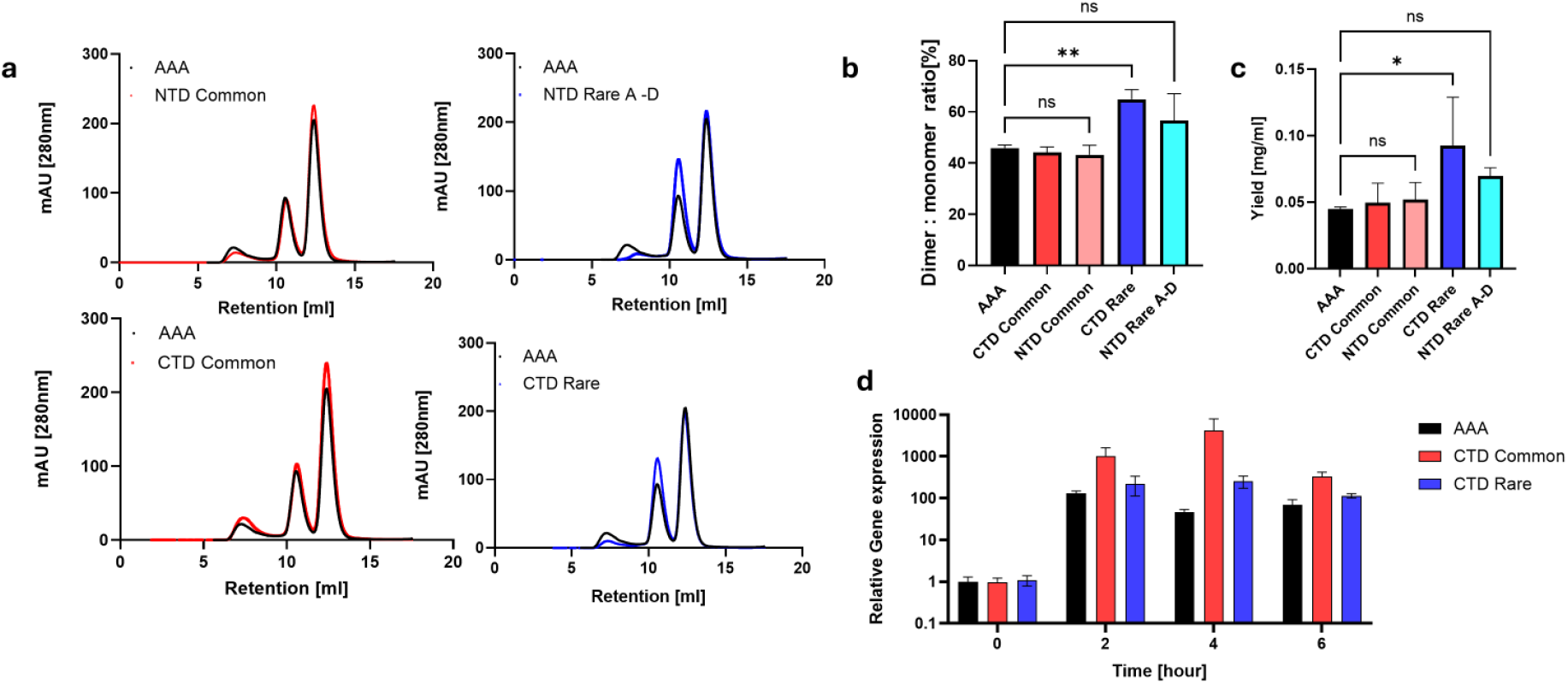
Synonymous codon usage modulates the dimerization propensityo in AAA myoglobin. (a) Size exclusion chromatography on synonymously coded AAA myoglobin demonstrated the that rare coding (blue) either before or after the hinge (top or bottom) variant and synonymously coded versions mutated to more abundant codons (NTD and CTD Common) or rarer codons (NTD and CTD Rare shades of blue) were analyzed on a SEC200 column to assess the monomer (last peak) – dimer (middle peak) ratio. (b) The calculated dimer to monomer ratio. (**p<0.0049 according to One-Way ANOVA) (c) Protein yield (mg/ml) per culture volume (\***p < 0.0288**, oneway ANOVA). (d) Relative gene expression levels.

Our findings demonstrate that synonymous coding can exert functional control over protein oligomerization by modulating the propensity for domain swapping. By altering the balance between monomeric and domain-swapped dimeric states in AAA myoglobin, our results provide direct experimental evidence that nucleotide-level sequence variation—without changes to amino-acid identity—can reshape protein assembly landscapes. In our system, rare coding yielded not only more domain swapped species, relative to the monomeric state, but also higher proportions of soluble protein and reduced the presence of higher-order soluble oligomeric states, consistent with improved co-translational folding and suppression of off-pathway assembly.

Synonymous mutations have increasingly been implicated in protein misfolding and aggregation by altering translational pausing, nascent chain exposure, and chaperone engagement, with documented consequences for proteostasis and cellular fitness (Walsh et al., 2020; Shen et al., 2022). In parallel, domain swapping has been widely proposed as a structural gateway to aggregation and amyloid formation, particularly when swapping becomes runaway or kinetically trapped (Bennett et al., 2006; Rousseau et al., 2011). By demonstrating that synonymous codon choice can alter domain-swapped dimerization, our work suggests a potential connection between codon usage and aggregation-relevant structural equilibria. More broadly, these findings support a model in which synonymous sequence variation contributes to disease risk not only through effects on expression or stability, but also by subtly reshaping folding and assembly pathways that determine whether proteins adopt functional or aggregation-prone states.

## Data Availability

The datasets created and analysed during the current study are available from the corresponding author on request.

## Methods

### Constructs

The horse heart myoglobin sequence (NCBI Reference Sequence: NM_001164016.2) was cloned into a pET28a vector between NdeI and XhoI as a thrombin clevable N-terminal construct (Twist Bioscience). The non synonymous mutant, AAA, was created by introducing hinge region H80A, H81A, G82A. Synonymous mutations were made to the AAA variant to create the constructs shown in Figure S3.

### Protein expression and Purification

A 10 ml starter culture was grown overnight in BL21 at 37°C and 200 rpm. The starter was diluted 1:100 t 1 L f LB c t 50 μ / L kanamycin and grown at 30°C (unless otherwise indicated for specific experiments shown in Figure S4), 00 u t l 600 ≈ 0 6–0.8 when expression was induced with 0.4 mM IPTG. The culture was further incubated at 20°C (except for the solubility screen experiments shown in Figure S4, where the incubation temperatures were screened as noted in the figure), 200 rpm overnight, and cells were harvested by centrifugation and either frozen at-20 °C or processed immediately for purification. For the solubility screen shown in Figure S4 the cells were lysed with BugBuster x10 protein extraction reagent (EMD Milipore Corp. USA) in 20mM Tris buffer pH 8. For all other experiments involving protein preparation, the cells were resuspended in lysis buffer (20mM Tris pH 8, 100mM NaCl) and disrupted by sonication. Following centrifugation, the lysate was loaded onto an equilibrated His Trap FF 5ml column (Cytiva) t fl t f 3 0 L/, th t ult v l t (UV)™v detector set at 280 nm. The elution buffer contained 20 mM Tris-HCl pH 8.0, 100 mM NaCl, and 300 mM imidazole. Protein purity was analyzed by15% SDS-PAGE, and protein concentrations were determined using a NanoDrop (BLUE-RAY BIOTECH EzDrop 1000). Finally, size-exclusion chromatography (SEC; Superdex 75 10/300 Cytiva) was conducted to analyze the purified protein Fractions. Fractions containing monomers or dimers were collected separately for downstream experiments.

### Gene Expression Assay

Starter cultures were grown overnight at 37 °C and 200 rpm. The cultures were diluted 1:50 t 50 L LB th 50 μ / L k yc t 30°C, 00 t 600 ≈ 0 6 when expression was induced with 0.4mM IPTG and the incubation temperature was reduced to 20 °C. Four hours post-induction, cultures were transferred to 50 ml centrifuge tubes and centrifuged at 5000 rpm for 20 minutes at 4 °C. The supernatant was discarded, and the bacterial pellet was resuspended in fresh 50 ml LB medium containing kanamycin. Samples were collected at 0, 2, 4, and 6 hours post-uct E ch l ‘600 ju t t 1 0 (≈10^9 c ll / L) Cells were treated with RNAprotect Reagent (Qiagen) according to the uf ctu’ t c l for subsequent RNA extraction. Total RNA was extracted with a commercial RNA isolation kit (MACHEREY-NAGEL). cDNA was synthesized using the Verso cDNA Synthesis Kit (ThermoFisher Scientific). Primer sequences were designed using Primer-BLAST (NCBI) (**Table S1**). qPCR was performed on the LightCycler 480 Instrument II (Roche) using 384-well plate format. The amplification program included one pre-incubation cycle at 95°C for 5 min, followed by 45 cycles of 95°C for 10s, 60°C for 20s, and 72°C for 20s. This was followed by a melting curve analysis cycle of 95°C for 5s and 65°C for 1 min. PCR amplification was performed with in 5 μl f ct xtu (L htCycl 480 YBR G II Master, Roche) containing a 1 /μL c NA and a final primer concentration of 250 nM. Expression was quantified relative to the mean expression of three housekeeping genes (ihfB, ldnT, hcaT) as reference genes (Zhou et al. 2011). Relative transcript abundance was c lcul t u th ΔC_T th (Yuan et al. 2006).

### Refolding profile of AAA protein using urea-induced denaturation

Monomeric and dimeric fractions of AAA, isolated via size-exclusion chromatography (Superdex 75 10/300 Cytiva), were exchanged into a buffer containing 8 M urea, 20 mM Tris-HCl (pH 8), and 100 mM NaCl using centrifugal filtration at 6000 rpm, 4 °C (10 kDa cutoff). Samples were further incubated at 4 °C for 3 h. Denatured samples were dialyzed against 20 mM Tris-HCl pH 8.0, 100 mM NaCl, 4°C overnight and then analyzed by size-exclusion chromatography (Superdex 75 10/300 Cytiva).

## Supplementary

**Figure S1:**
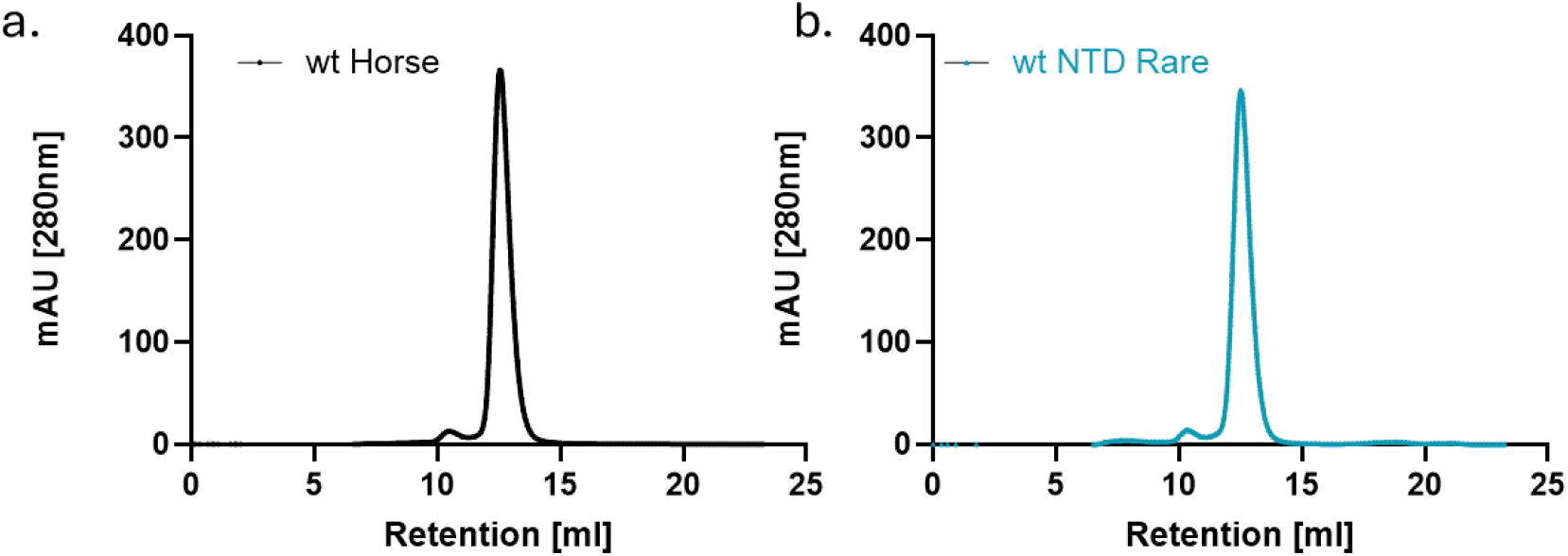
wt horse heart myoglobin forms a small fraction of domain swapped dimer. (a) size exclusion chromatogram of 4mg/ml wt Horse heart myoglobin. Retention time 10.49 ml presents the domain-swapped dimer fraction, whereas the 12.55 ml fraction represents the monomer fraction. (b) size exclusion chromatogram of 4mg/ml wt Horse heart myoglobin with rare codon usage (wt NTD Rare). Retention time 10.35 ml presents the domain-swapped dimer fraction, whereas the 12.52 ml fraction represents the monomer fraction.

**Figure S2.**
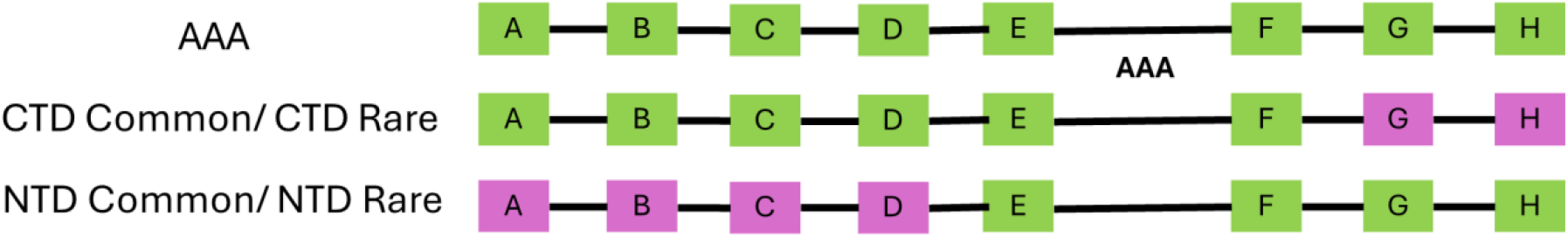
Mapping of synonymous mutation to protein regions. Horse heart myoglobin has eight helices (labelled A to H), with the domain swapping hinge located between helix E and F. N terminal domain (NTD) mutations were made across the sequence spanning helices A to D. C terminal domain (CTD) mutations were made across the sequence spanning helices G and H.

**Figure S3.**
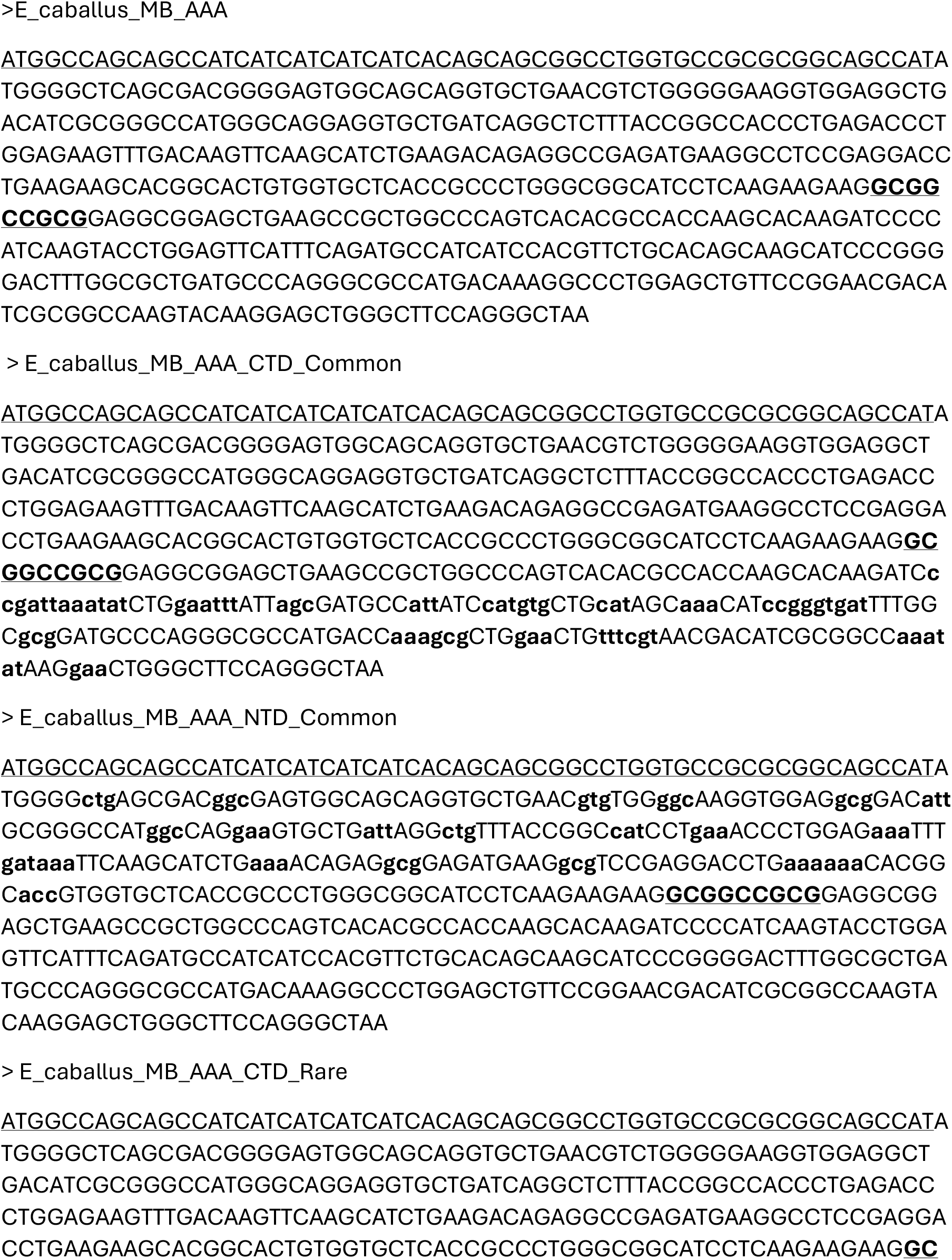
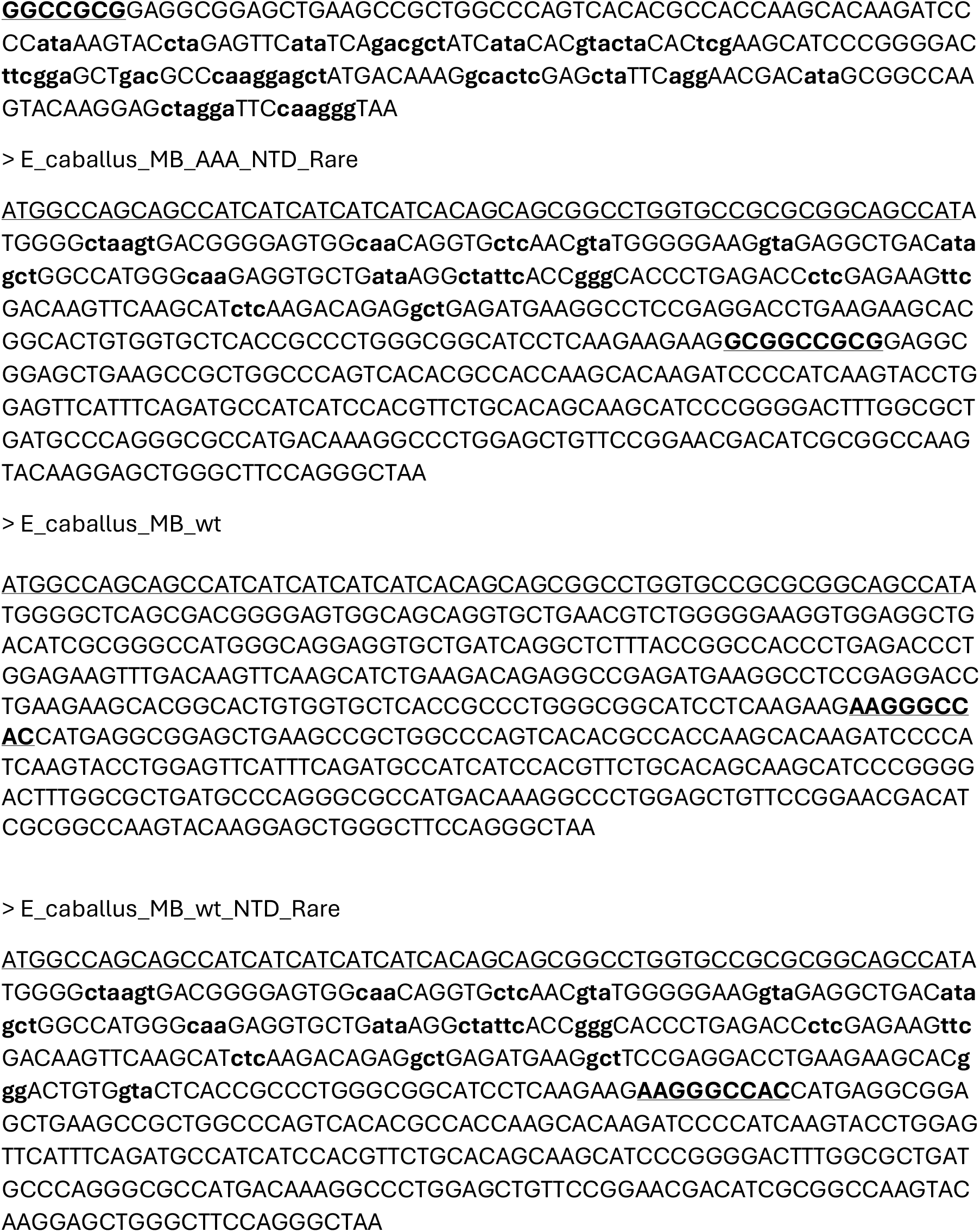
Fasta sequences of genes used in this study. Synonymous mutations from the wt gene sequence are highlighted with bolded lower case letters. The hinge region is highlighted with bolded underlined letters. The coding sequence of the N terminal thrombin cleavable His tag is highlighted by underlining only.

**Figure S4.**
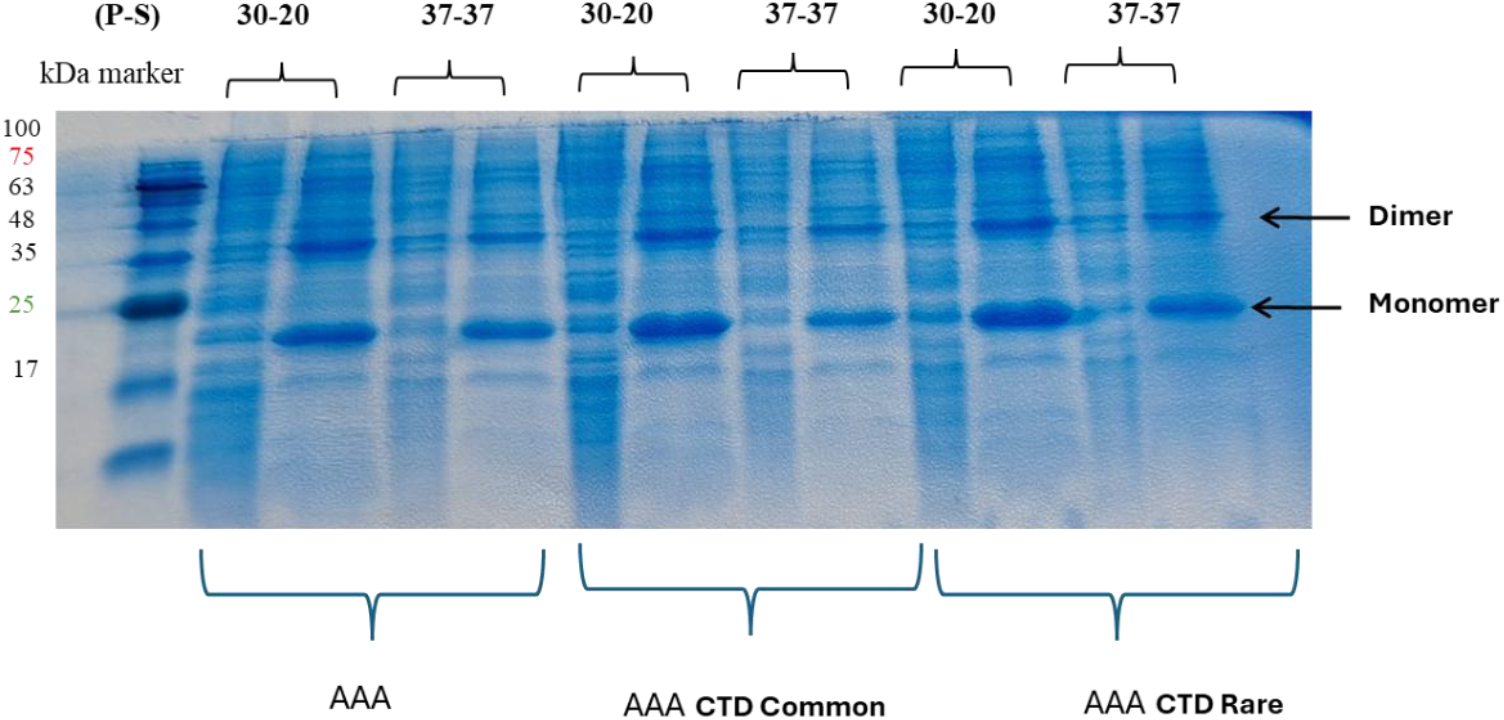
Optimal expression temperature scan for AAA variants. The effect of pre and post induction temperature on protein solubility was screened across synonymous mutants by visualizing the non-soluble fraction (P) and soluble (S) fraction on a 15% SDS PAGE gel. Across all conditions, and all variants tested, both monomeric and dimeric species were observed and, in all cases, both oligomeric states of the protein was observed to be mostly soluble.

**Table S1:**
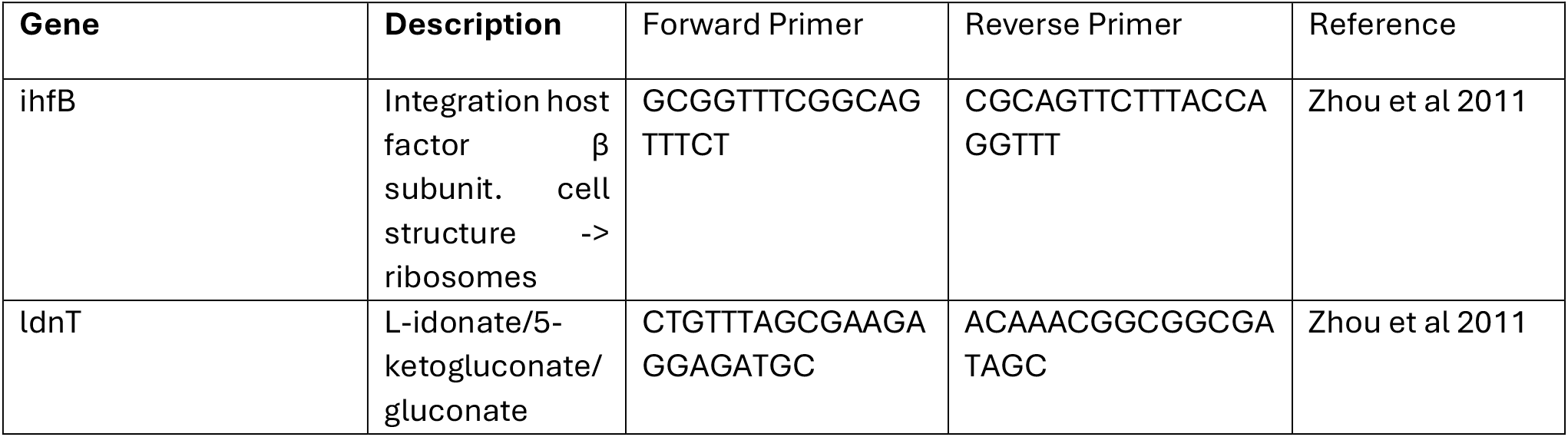
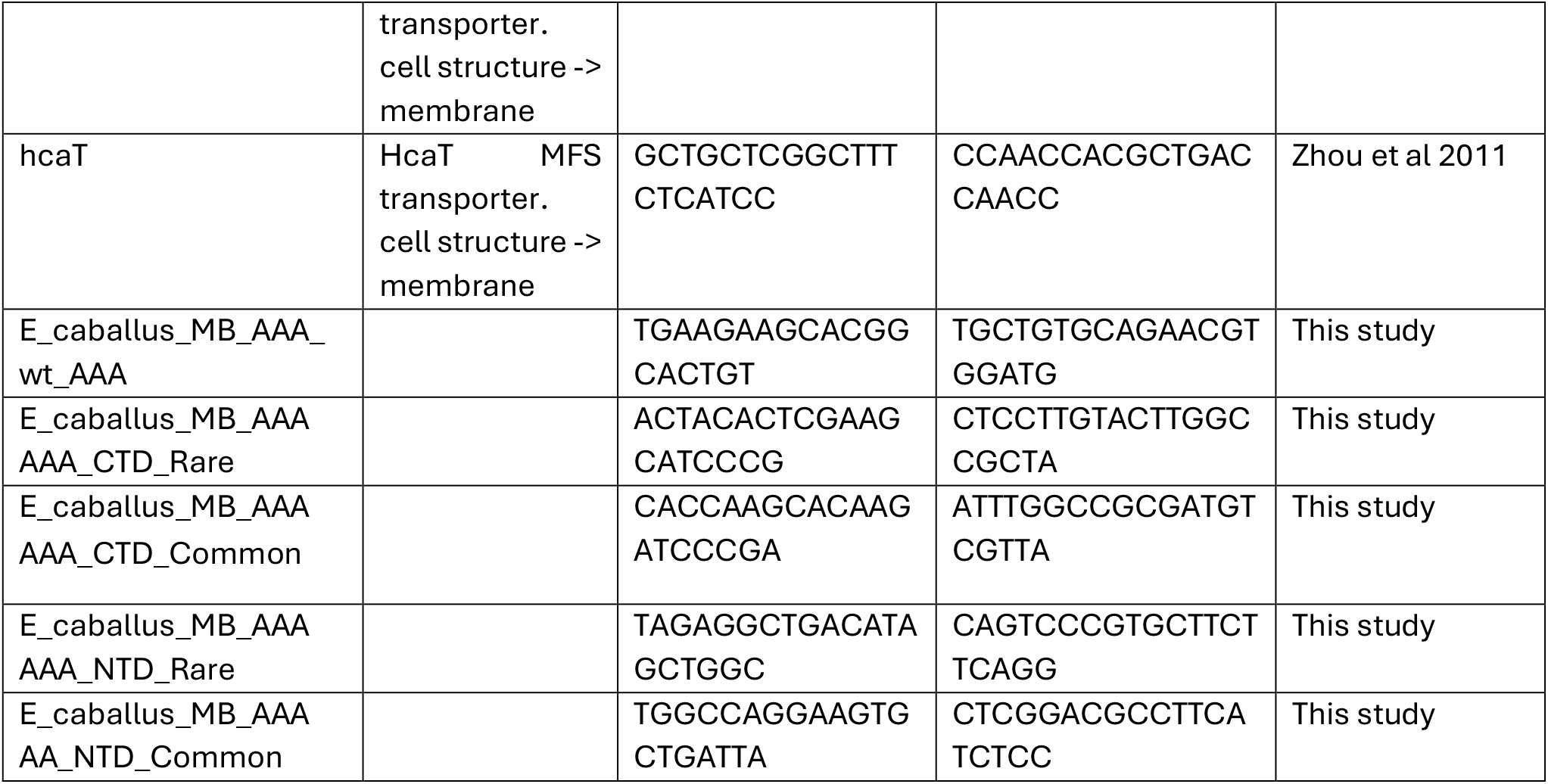
Gene Expression Assay Primers.

